# Comparing the prioritisation of items and feature-dimensions in visual working memory

**DOI:** 10.1101/863191

**Authors:** Jasper E. Hajonides, Freek van Ede, Mark G. Stokes, Anna C. Nobre

## Abstract

Selective attention can be directed not only to external sensory inputs, but also to internal sensory representations held within visual working memory (VWM). To date, this has been studied predominantly following retrospective cues directing attention to particular items, or their locations in memory. In addition to item-level attentional prioritisation, recent studies have shown that selectively attending to feature dimensions in VWM can also improve memory recall performance. However, no study to date has directly compared item-based and feature-based attention in VWM, nor their neural bases. Here, we compared the benefits of retrospective cues (retro-cues) that were directed either at a multi-feature item or at a feature-dimension that was shared between two spatially segregated items. Behavioural results revealed qualitatively similar attentional benefits in both recall accuracy and response time, but also showed that cueing benefits were larger following item cues. Concurrent EEG measurements further revealed a similar attenuation of posterior alpha oscillations following both item and feature retro-cues when compared to non-informative, neutral retro-cues. We argue that attention can act flexibly to prioritise the most relevant information – at either the item or the feature-level – to optimise ensuing memory-based task performance, and we discuss the implications of the observed commonalities and differences between item-level and feature-level prioritisation in VWM.

## Introduction

Visual working memory (VWM) provides a means to maintain relevant information independently of continued visual input, to guide adaptive behaviour (Baddeley, 1992, 2003). Because VWM has limited capacity and/or resources (Bays & Husain, 2008; Luck & Vogel, 1997; Vogel, Woodman, & Luck, 2001; Zhang & Luck, 2008), it is essential to distribute memory processes efficiently to complete tasks at hand effectively. Over the past decade, it has become increasingly clear that VWM is more flexible than originally thought. Focused attention continues to prioritise and select contents maintained in VWM as goals and predictions about goals change (Griffin & Nobre, 2003; Kuo, Stokes, & Nobre, 2011; Landman, Spekreijse, & Lamme, 2003; Souza & Oberauer, 2016). To bring about behavioural benefits, attention-related modulatory signals must interact with mnemonic information that is available within VWM. Thus, by studying what forms of attention confer benefits to VWM also provides insight about the format of information held in VWM.

In perceptual attention, research has revealed a multitude of representational formats available for modulation. These include spatial locations, objects, features, semantic associations, time intervals, and likely more (for an overview, see e.g., Nobre, 2018). Whether the same diversity of representational formats is available in VWM is an important and informative question. Information in VWM results from attentional filtering of incoming sensory processing (Vogel, McCollough, & Machizawa, 2005). Thus, the representational information might be kept in an altered, more compact format. For example, it has been suggested that the primary representational unit of VWM involves integrated (feature-bound) items (Luck & Vogel, 1997; Vogel et al., 2001). Under this framework, one might expect that the primary target for attentional selection in visual working memory should be at the level of individual items. Accordingly, most studies looking at the role of attention in VWM to date have used spatial retro-cues, linked to item-based representations, and have shown clear benefits (e.g. Griffin & Nobre, 2003; Landman et al., 2003; Souza & Oberauer, 2016).

At the same time, recent studies have demonstrated that attention can also facilitate behaviour when directed to *feature dimensions* that are shared among multiple items in VWM (Niklaus, Nobre, & Van Ede, 2017; Park, Sy, Hong, & Tong, 2017; Pilling & Barrett, 2016; Ye, Hu, Ristaniemi, Gendron, & Liu, 2016; Yu & Shim, 2017). It remains unclear, however, how such effects of feature-based attention compare to item-based attention in VWM, as no study has directly compared these two forms of attentional facilitation in VWM. Here we directly compare these two types of attentional facilitation.

In addition to comparing item and feature-based attention at the level of behavioural performance, we examined their effects on an electrophysiological marker linked to attention in VWM: the attenuation of posterior alpha oscillations. Several studies have revealed that item-based prioritisation in VWM is associated with the attenuation of alpha oscillations in posterior brain areas, suggesting modulation of visual areas involved in representing the mnemonic items (Myers, Walther, Wallis, Stokes, & Nobre, 2015; Poch, Capilla, Hinojosa, & Campo, 2017; van Ede, 2018; van Ede, Niklaus, & Nobre, 2017; Wallis, Stokes, Cousijn, Woolrich, & Nobre, 2015; Wolff, Jochim, Akyürek, & Stokes, 2017). It remains unclear whether alpha attenuation also occurs during the attentional prioritisation of feature dimensions that are shared across multiple items held in VWM.

In the current study, we therefore compared and contrasted behavioural and neural effects of internal shifts of attention to multi-feature items and to single feature dimensions that were shared across multiple items. Through the behavioural data, the aim was to test whether there is a clear primacy of the object-level information in VWM. If the representational format in VWM organises items as integrated objects, this should also be the primary level at which attention can operate. Accordingly, benefits from item-directing retro-cues should be substantially larger. If, however, attention has similar access to multiple levels of information in VWM, then retro-cueing benefits for feature dimensions and individual items may be similar. By recording EEG and measuring alpha oscillations, we further tested whether a similar alpha modulation occurs when attention is directed to a cued item or to a visual feature dimension that is distributed across multiple items held in VWM.

To address these questions, we used a task in which participants were presented with two Gabor gratings, each of which contained both colour and orientation information. On half of the blocks, participants were presented with an item-directing retro-cue and on the other half with a feature-dimension-directing retro-cue. Both blocks contained neutral (uninformative) retro-cues, against which we compared the effects of both types of informative retro-cues. We observed qualitatively comparable retro-cueing effects, though benefits following item cues were larger. Both effects were accompanied by similar alpha attenuation following the cues, and both item- and feature-level benefits on behaviour were highly correlated across participants.

## Methods

### Participants

The study was approved by the Central University Research Ethics Committee of the University of Oxford. Thirty-two healthy volunteers (19 female; mean age 28.3; range 18-35) took part. Participants had normal or corrected-to-normal vision and were not colour blind. Participants provided written informed consent before participating in the study and were paid £15 per hour. Data from two participants were excluded from analysis, one for terminating the experiment early and the other due to hardware failure.

### Experimental set-up & stimuli

Participants were seated in front of a 23-inch monitor (1920 × 1080, 100 Hz). Stimuli were generated using Psychophysics Toolbox version 3.0.11 (Brainard, 1997) in MATLAB 2014b (MathWorks, Natick, MA). Head position was set at 90 cm from the monitor, and participants used a chinrest. The stimuli consisted of luminance-defined sinusoidal Gabor gratings generated in MATLAB 2014b. Fourty-eight evenly spaced colours were drawn from a circle in CIE L*a*b colour space (center at L = 54, a = 18, b = −8, radius =59). Gratings were presented using one of 48 different orientations (3.75 to 180 degrees in steps of 3.75) and 48 different colours.

### Task & design

Participants performed a visual working memory task (**Figure 1**) in which they were asked to reproduce the colour or orientation of one out of two memory items at the end of a memory delay of 2.3 seconds. At the start of each trial, two Gabor stimuli with a radius of 2.2 degrees positioned left and right from fixation (centred 3.1 degrees of visual angle) were presented simultaneously for 300 ms. Participants were instructed to remember the colour and the orientation of both items. At the end of the trial, they were probed to report the orientation or the colour of one of the items. The to-be-reported feature was indicated with the probe circle that was either a colour wheel (colour report) or a white wheel (orientation report), while the to-be-reported item was indicated by the location of the probe circle (left/right, corresponding to the original location of the probed memory item). Orientation and colour values varied independently between the two items, with the constraint that no two equal orientations or colours were presented on the same trial. Colours and orientations were counterbalanced so that each was presented equally often across trials.

**Figure 1.**
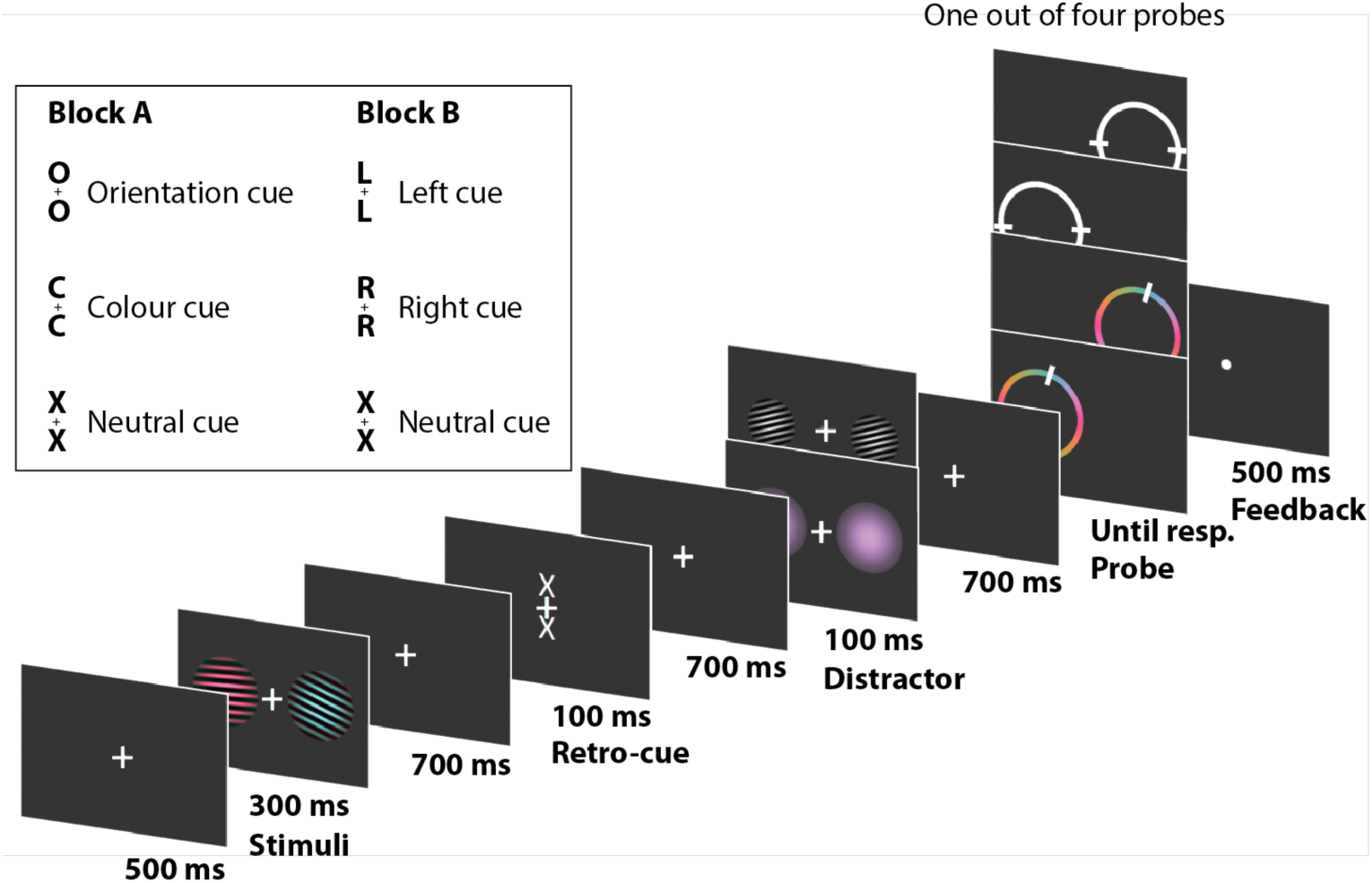
Experimental design with timings. Participants were presented with two coloured Gabor gratings to memorise. Subsequently, a retro-cue informed participant which item or feature dimension would be probed. After either a colour distractor or an orientation distractor with a semi-randomly drawn orientation or feature, the probe was presented. Participants adjusted the probe dial to match the feature in memory.

Two events occurred during the memory delay: first a retro-cue appeared, which could provide information about the item or feature dimension that would be probed. Second, 700 ms after the retro-cue offset, an irrelevant ‘distractor’ stimulus was presented that contained either colour or orientation information (**Figure 1** for examples).

On even runs, informative retro-cues indicated the location (left, ‘L’, or right, ‘R’) of the item that would be probed at the end of the trial, without giving information about what feature would be probed. On odd runs, informative retro-cues indicated the feature dimension to be probed (colour, ‘C’, or orientation, ‘O’), without giving information about what item would be probed. We thus cued either a single item that contained two relevant features, or a single feature-dimension that was shared between two relevant items. When informative, the retro-cue was always 100% valid. Both runs also contained 50% non-informative neutral cues (‘X’) that provided no information about what item or feature would be probed at the end of the trial. Participants were encouraged to use the informative retro-cues to select the relevant item or feature dimension.

Following another fixation period of 700 ms (after the retro-cue), a bilateral distractor was presented for 100 ms. The distractor consisted of either a single colour or a black-and-white oriented Gabor grating. Sixteen colours and 16 orientations were used (11.25 to 180 degrees in steps of 11.25 degrees). These varied randomly from the orientation/colour of the items in the memory array. However, we ensured that the three types of cues (C/O/X or L/R/X) all contained the same range of 16 distractor features. Out of these 16 distractor features, we randomly assigned 8 as colour features and 8 as orientation features.

After another fixation period of 700 ms (after the distractor), the colour-wheel or a white-circle probe appeared. To keep the orientation and colour recall as similar as possible, we presented the colours at a fixed position on the colour wheel. Participants were instructed to respond as accurately as possible by using the ‘J’ and ‘F’ key to rotate the probe counter-clockwise and clockwise, respectively. Participants were instructed to use their left index finger to press the ‘J’ key and the right index finger to press the ‘F’ key. Although there was no explicit time limit for the response time, we logged reaction times as the time between the onset of the probe and the first button press that initiated the ‘dial-up’ report. Reaction time therefore serves as a proxy for the time it took participants to access the relevant memory information before commencing their reproduction report.

### Behavioural analysis

We computed the error for each trial for each participant by subtracting the target orientation or colour (in radians around the colour circle in CIE L*a*B space) from the probe response. All error scores were mapped onto a −½π to ½π space. All trials for which the reaction time was more than 4 standard deviations above a participant’s mean decision time were discarded (0.9% ± 0.3%). To calculate the retro-cue benefit, we subtracted the absolute error on cued trials from the absolute error on neutral trials. In all our analyses, we only compared trials of one retro-cueing condition with neutral trials from the same blocks.

A mixture model was fitted separately for each retro-cueing condition and respective neutral condition, modelling target response rate, guess rate, swap responses, and precision to the error data of each subject (Bays, Catalao, & Husain, 2009; Zhang & Luck, 2008). We fitted the mixture model separately for every subject, colour or orientation recall, spatial or feature retro-cues, and informative or neutral retro-cues. Estimating the mixture-model parameters allowed for estimation of different components that contribute the overall error; we estimated the fidelity of the representation independently of the guess and swap rate. We used the mixture model made available by Bays et al. (2009).

When comparing more than two conditions, we applied a repeated-measures analysis of variance (rmANOVA) and report η^2^ as a measure of effect size. When evaluating retro-cueing benefits, we applied dependent samples t-test, comparing informative vs neutral cues, as well as the cueing effects between item and feature retro-cues. When the assumption of normality was violated we instead applied a Wilcoxon signed-rank test. We report Cohen’s d as a measure of effect size for parametric tests and matched rank biserial correlation for non-parametric effect size. For evaluation we two-sided tests with a critical alpha value of 0.05.

### EEG acquisition

EEG data were collected using Synamps amplifiers and Neuroscan software (Compumedics). We used a 61 Ag/AgCl sintered electrodes (EasyCap, Herrsching, Germany), laid out according to the international 10-10 system, with mastoids behind the left and right ear. The left mastoid was used as an active reference during the recordings. Offline, an average-mastoids reference was derived using the left and right mastoids. The ground electrode was placed on the left arm above the elbow. Horizontal EOG was measured using lateral electrodes next to both eyes while vertical EOG was measured above and below the left eye. Data were sampled at 1000 Hz, and stored for subsequent analysis.

### EEG preprocessing

Data were imported into MATLAB 2017a using *pop_loadcurry()* and further analysed using Fieldtrip (Oostenveld, Fries, Maris, & Schoffelen, 2011) and the OHBA Software Library (OSL; https://ohba-analysis.github.io/). Analysis started by cutting out the epochs between 100 ms before and 2200 ms after retro-cue onset (ft_redefinetrial) followed by re-referencing the data to the average of the mastoids (ft_preprocessing). EEG data were down sampled to 200 Hz to reduce computational demands and storage space (ft_resampledata).

Next, EEG data were further de-noised using Independent Component Analysis (ICA; ft_componentalanalysis) applying the FastICA algorithm (Hyvärinen, 1999) to all EEG sensors. ICA separates the EEG signal into non-Gaussian subcomponents of the data that are statistically independent from one another. Spatial components strongly correlated (r > 0.4) with electrooculogram (EOG) channels were removed from the EEG data. We set-out to remove trials on which participants blinked during the window of 100 ms prior up to 200 ms post retro-cue presentation. After baselining the horizontal EOG signal at −300 to −100 ms trials on which horizontal EOG voltage surpassed 200 μV (approximately ½ of the maximum voltage evoked by a typical blink) were flagged and later removed from EEG and behavioural analyses (0.446% ± 1.23%; mean ± standard deviation). Subsequently, we removed epochs based on within-trial variance of the broadband signal at a 0.05 significance threshold using a generalised ESD test (Rosner, 1983; implemented in OSL) and discarded 2.48% ± 2.18% (mean ± standard deviation) of the trials.

### Time-frequency processing

Time-frequency decomposition of the EEG signal was done using ft_freqanalysis. Spectral power between 2 and 50 Hz was computed on Hanning-tapered data using a short-time Fourier Transform, with a 300-ms sliding time window that was advanced in steps of 15 ms. We zoom in on modulations in posterior alpha oscillations, by averaging the time-frequency plots for the 17 most posterior electrodes and calculating the normalised differences in power following between informative and neutral retro-cues ([informative – neutral]/[informative + neutral] × 100). We did this separately for left and right item retro-cues, and for colour and orientation feature retro-cues. We also compared left vs. right, and colour vs. orientation, retro-cue conditions directly using the same quantification. For statistical evaluation, we applied a two-sided cluster-based permutation analysis (Maris & Oostenveld, 2007) with 5000 permutations at an evaluation threshold of 0.05.

To characterise the onset of alpha attenuation after the retro-cue, we extracted the time course of 7-12 Hz power modulation (in the specified informative vs. neutral cue contrast) and focused on the 0-1000 ms period post retro-cue onset. On these data, we then identified the earliest timepoint in which the power modulation reached half of its minimal value for each condition. This latency was used as a measure to compare neural modulation by feature retro-cues and item retro-cues.

To depict the topography of the power modulations analysed in the predefined set of posterior electrodes (depicted in **Figure 4**), we calculated the relevant contrast for each electrode and averaged over the time-frequency window of 400-800 ms and 7-12 Hz for the values of all posterior electrodes (O2, PO8, PO4, P8, P6, P4, P2, O1, PO7, PO3, P7, P5, P3, P1, Oz, POz, Pz). In addition, to focus on alpha lateralisation, we contrasted activity in electrodes left posterior electrodes (O1, PO7, PO3, P7, P5, P3, P1) and right posterior electrodes (O2, PO8, PO4, P8, P6, P4, P2), contralateral vs ipsilateral to the cued item following informative item retro-cues.

**Figure 2.**
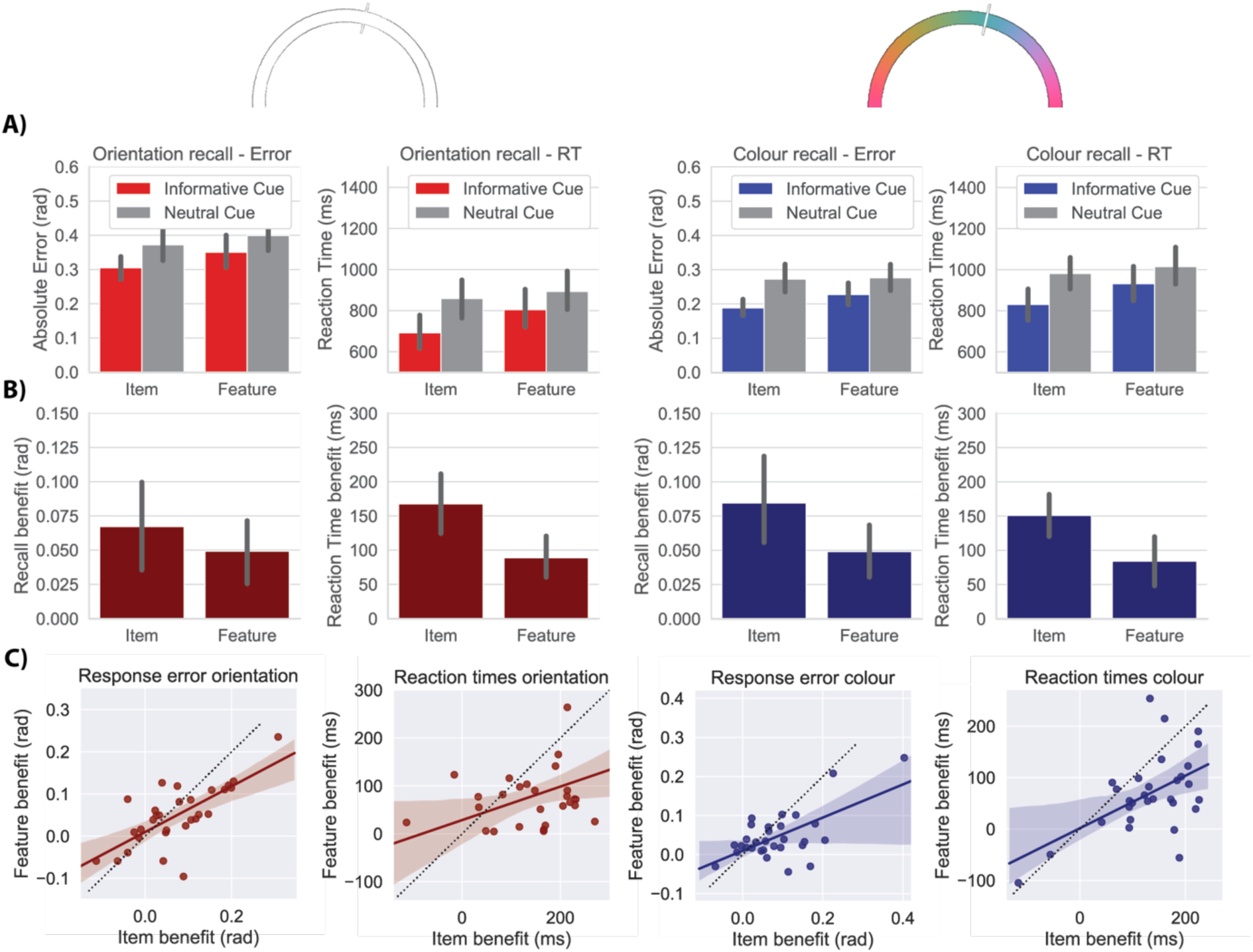
Performance benefits of item and feature-dimension retro cueing. A) The four panels show absolute error and reaction times for trials with an informative cue or a neutral cue. Trials in which orientation was probed are displayed in red while colour-probe trials are displayed in blue. Each panel shows the data separately for item retro-cue blocks and feature retro-cue blocks. B) Behavioural benefit of retro-cues. Subtracting the mean absolute error on trials with an informative cue from the neutral trials gives the performance benefit of the retro-cue – here expressed as positive values. Orientation benefit is depicted in dark red and colour benefit in dark blue. C) Correlations across participants between the item and feature retro-cue benefits in shown in B. Error bars show 95% confidence intervals.

**Figure 3.**
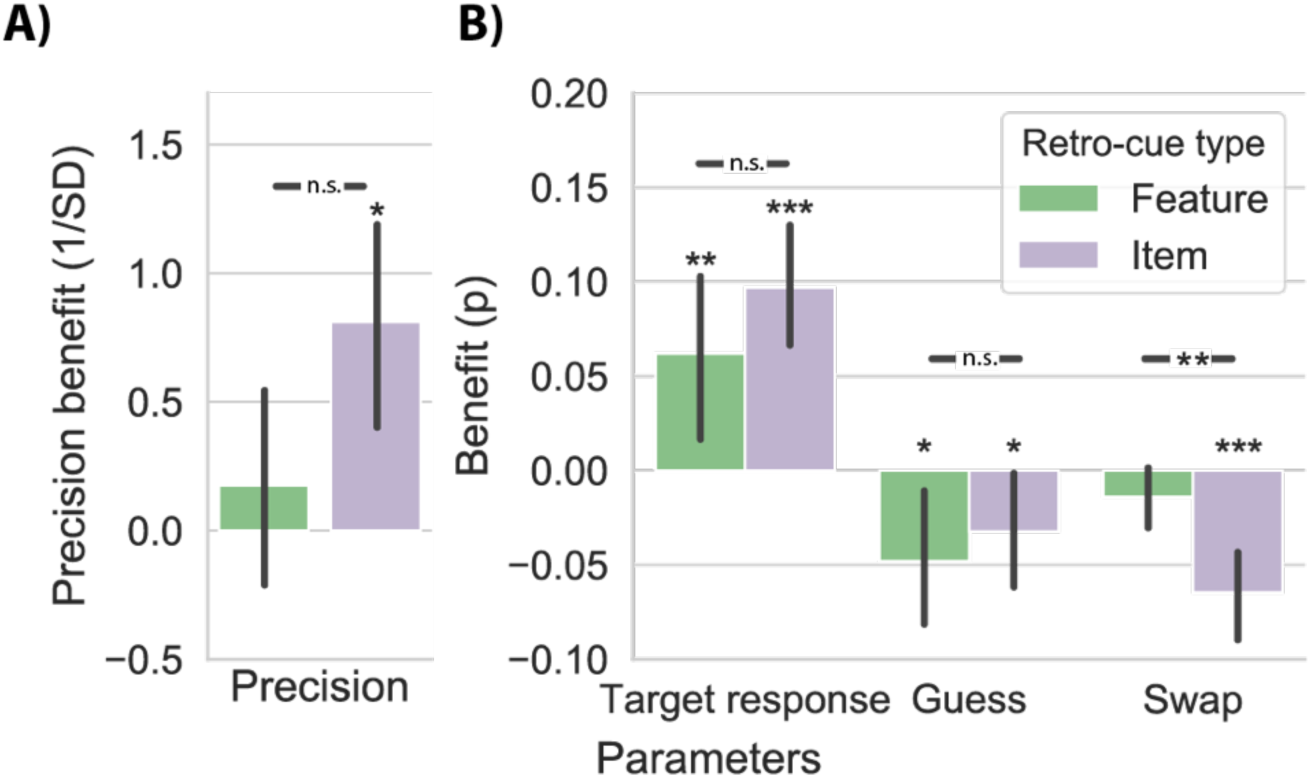
Cueing effects on mixture modelling parameters. A) Mixture-model estimates for the cueing effects on A) precision, B) target response, guess rate, and swap rate, for trials where item retro-cues or feature retro-cues were presented compared to neutral trials. Hence, valid retro-cues positively influenced target response proportions and negatively influenced guess rate and swap rate. The green and purple asterisks indicate significant differences of, respectively, item- or feature benefits from zero (i.e., benefits following informative vs. neutral cues). Black asterisks indicate significant differences between item and feature retro-cueing benefits. Error bars indicate 95% confidence intervals. * p < .05, ** p < .01, *** p < .001.

**Figure 4.**
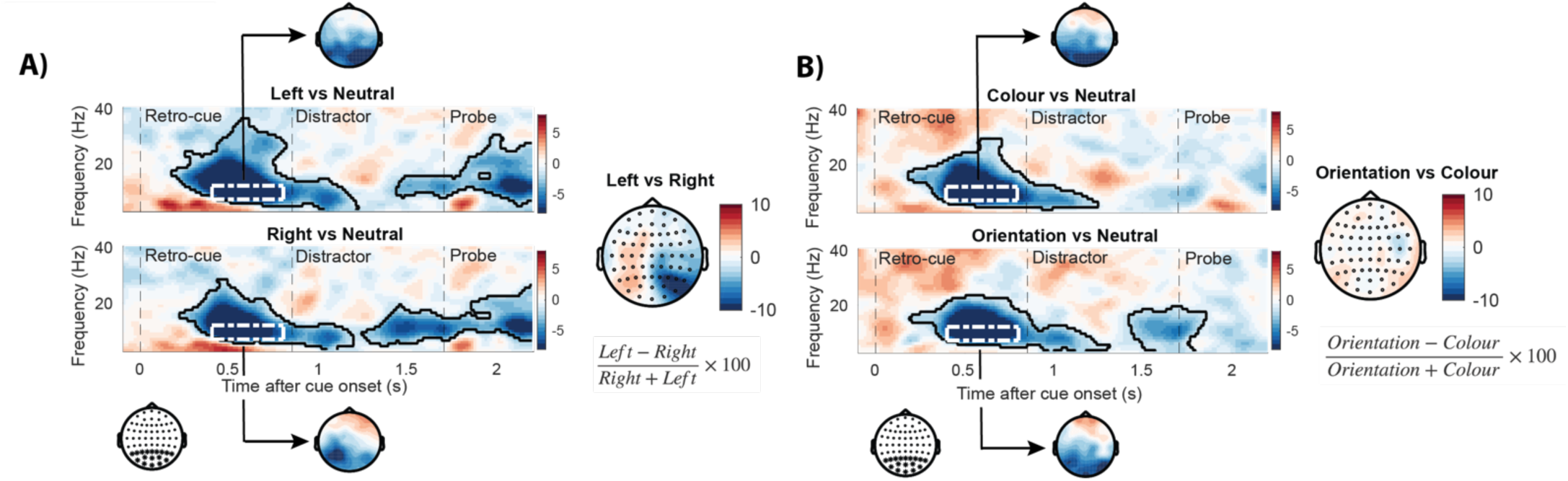
Induced neural EEG responses to the left and right item retro-cues and to colour and orientation feature retro-cues. A) Time-frequency representation of the difference between left/right cued trials vs. neutral trials in item-cue blocks in the predefined posterior electrode cluster indicated with asterisks below the plots. The topography to the right shows the difference between left and right retro-cues (**Supplementary Fig. 2** for the corresponding time-frequency map of the difference between contralateral and ipsilateral responses). B) Same representations as outlined above but here we compare colour and orientation with their respective neutral trials or with one another. Highlighted areas with the black solid outline indicate significant clusters (permutation test, n = 30, cluster-forming threshold p < .05, corrected significance threshold p < .05). The topographies display the alpha power (7-12 Hz) in the 400-800 ms window that is also demarcated in the time-frequency plots with the white dashed boxes.

Topographies were intended solely to portray the nature of the modulation and were not subjected to further statistical testing.

## Results

**Figure 2A** shows behavioural performance as a function of experimental condition (collapsed over distractor type, as this did not yield consistent results as discussed below). To analyse the effects of item and feature-dimension retro-cues, we quantified retro-cueing benefits as the difference between the trials with informative and neutral retro-cues (**Figure 2B**).

To quantify formally the effects of retro-cue informativeness (valid or neutral) and retro-cue block type (item retro-cue block or feature retro-cue block) we used a 2 × 2 rmANOVA. We ran this separately for RT and response error, and separately for both colour and orientation recall reports. We observed a significant main effect of retro-cue informativeness, with better performance following informative vs. neutral retro-cues on all four dependent variables: orientation error (*F*_1,29_ = 18.888; *p* < .001; η^2^ = 0.394), colour error (*F*_1,29_ = 26.387; *p* < .001; η^2^ = 0.476), orientation RT (*F*_1,29_ = 67.27; *p* < .001; η^2^ = 0.699), and colour RT (*F*_1,29_ = 65.15; *p* < .001; η^2^ = 0.692). At the same time, we found that the behavioural benefits of retro-cue informativeness were larger in item retro-cue blocks than in feature retro-cue blocks, yielding a significant interaction for colour error (*F*_1,29_ = 9.065; *p* = .005; η^2^ = 0.238), orientation RT (*F*_1,29_ = 19.64; *p* < .001; η^2^ = 0.404), and colour RT (*F*_1,29_ = 21.00; *p* < .001; η^2^ = 0.420). Though we found the same trend, this did not reach significance for orientation error (*F*_1,29_ = 3.750; *p* = .063; η^2^ = 0.115). Finally, in line with the larger benefit of item-retro-cues, we also found a significant main effect of block type, constituted by better performance in item retro-cue blocks for all four dependent variables: orientation error (*F*_1,29_ = 39.634; *p* < .001; η^2^ = 0.577), colour error, *F*_1,29_ = 8.343; *p* = .007; η^2^ = 0.223),orientation RT, *F*_1,29_ = 50.51; *p* < .001; η^2^ = 0.635), and colour RT (*F*_1,29_ = 32.08; *p* < .001; η^2^ = 0.525).

For completeness, we also considered the third factor ‘distractor congruence’ (i.e., when the distractor contained the same or the other feature dimension as the to-be-recalled memory feature), but found no systematic effects of distractor congruence across our four dependent variables, nor interactions with the factors of interest – see **supplementary table 1**.

In the following, we describe in more detail the item- and feature retro-cueing effects of interest, in accordance with the data presented in **Figure 2**.

For orientation recall reports, participants significantly benefitted from item retro-cues. They had smaller errors (*t*_29_ = 4.235; *p* < .001; d = 0.773) and responded faster (*t*_29_ = 7.854; *p* < .001; d = 1.434), compared to trials with neutral retro-cues in the same blocks. Similarly, orientation reports benefitted significantly from feature cues in both reproduction error (*t*_29_ = 3.748; *p* = .001; d = 0.684) and response onset time (*t*_29_ = 6.302; *p* < .001; d = 1.151) compared to neutral trials within the feature retro-cueing blocks. Item cues conferred numerically larger benefits than feature cues. The difference was not statistically significant for error (0.023 radian, 48%; *t*_29_ = 1.936; *p* = .063; d = .354), but reached significance for reaction times (79 ms, 94%; *t*_29_ = 4.431; *p* < .001; d = 0.809).

The same pattern of results was found for the error and reaction times in the colour recall trials: colour reports benefitted from both item cues (*t*_29_ = 5.060; *p* < .001; d = 0.924) and feature cues (*t*_29_ = 4.069; *p* < .001; d = 0.743) and responses were also faster for item cues (*t*_29_ = 9.097; *p* < .001; d = 1.661) and feature cues (*t*_29_ = 4.951; *p* < .001; d = 0.904) compared to their respective neutral trials. For colour reports, we also found greater benefits of item retro-cues compared to feature retro-cues for both error (0.039 rad, 86%; *t*_29_ = 3.011; *p* = .005; d = 0.550) and reaction time (63 ms, 91%; *t*_29_ = 4.583; *p* < .001; d = 0.837).

Benefits of item-based and feature-based retro-cueing showed strong positive correlations across individuals for both colour and orientation reports (**Figure 2C**). For orientation reports, we found significant correlations between retro-cueing benefits following item cues and feature cues for both error (*r* = .709; *p* < .001) and reaction time (*r* = .539; *p* = .002). Likewise, for colour reports, we found significant correlations between retro-cueing benefits following item cues and feature cues for both error (*r* = .697; *p* < .001) and reaction time (*r* = .596; *p* < .001). Thus, participants who benefitted most from item retro-cues also benefitted most from feature retro-cues.

### Mixture modelling

In addition to the raw behavioural scores, we also modelled sources of error using a mixture model (**Figure 3AB**; Bays et al., 2009). We modelled four components 1) precision, characterised by width (1/STD) of the target centred response distribution, 2) proportion of target responses modelled by the gaussian centred around the target, 3) proportion of random responses characterised by the height of the uniform response distribution, 4) proportion of responses to the non-cued feature of the same dimension as the cued feature (non-target report or ‘swap’ errors). **Figure 3A** and **B** show the retro-cueing effects (informative vs. neutral) on each of these four parameters, separately for item and feature-cues (collapsed over colour and orientation reports, after fitting the model for each condition separately; see **Supplementary figure 1** for mixture model parameters separated for colour and orientation reports). As depicted in **Figure 3A,B** informative (vs. neutral) retro-cues significantly increased precision for item retro-cues (item: *t*_29_ = 2.736; *p* = .011; d = 0.500) though this did not reach significance for feature retro-cues: *t*_29_ = 0.578; *p* = .568; d = 0.105). At the same time, both item and feature retro-cues increased target response rates (item: *t*_29_ = 5.595; *p* < .001; d = 1.022; feature: Z_29_ = 382; *p* = .001; r_rb_ = 0.643), and decreased guess rates (item: *t*_29_ = −2.131; *p* = .042; d = −0.389; feature: Z_29_ = 78; *p* < .001; r_rb_ = −0.665), and item retro-cues further decreased swap rate (item: Z_29_ = 21; *p* < .001; r_rb_ = −0.910; feature: Z_29_ = 137; *p* = .080; r_rb_ = −0.368).

Direct comparisons between item and feature retro-cue benefits showed a significantly greater reduction in the rate of swap errors by item retro-cues relative to feature retro-cues (Wilcoxon signed-rank test; Z_29_ = 95; *p* = 0.002; r_rb_ = 0.591; see **Figure 3AB**). Effects for the other three parameters were not statistically different between item and feature retro-cues (all p > .10).

### Alpha attenuation following feature and item retro-cues

**Figure 4** shows the time- and frequency-resolved modulations in spectral EEG power in posterior electrodes following item and feature retro-cues, expressed as a difference from the neutral retro-cueing condition (neutral retro-cue minus informative retro-cue). After both item retro-cues and feature retro-cues, we observe an attenuation of alpha power starting at around 400 ms after presentation of the retro-cue (clusters all conditions p < .001). The alpha attenuation in trials with informative retro-cues re-emerges after the distractor onset, in the window just prior to the probe. To reveal the spatial layout of the significant clusters, we visualised the EEG topographies of the alpha-band power at 400 to 800 ms after the retro-cue. Similar topographies were associated with the later alpha modulation after the distractor and with the early modulation in the higher 13-30 Hz band (topographies not depicted).

In addition to this ‘global effect’ when comparing informative to neutral retro-cues, we also evaluated the difference between left and right item cues, and between colour and orientation feature cues (**Figure 4AB** right topographies, see also **Supplementary Figure 2** for corresponding time-frequency maps). In line with several prior studies (Myers et al., 2015; Poch et al., 2017; van Ede et al., 2017; Wallis et al., 2015; Wolff et al., 2017), following item cues, alpha attenuation was most pronounced contralateral to the memorised location of the cued item. In contrast, following feature retro-cues no clear differences were observed between colour and orientation cues, which directed attention to a single feature-dimension that was shared between the left and right items. Finally, we found that the alpha attenuation had very similar latencies following item-directing and feature-directing retro-cues (t_29_ = 1.273; *p* = .213; **Supplementary Figure 3**).

## Discussion

We demonstrate that both item-based and feature-based attentional prioritisation during VWM maintenance decreases recall error and speeds response initiation times following the probe. Hence, we replicate the finding that selective attention can retrospectively prioritise not only items (Griffin & Nobre, 2003; Kuo et al., 2011; Landman et al., 2003; Souza & Oberauer, 2016), but also feature dimensions maintained in VWM (Niklaus et al., 2017; Park et al., 2017; Ye et al., 2016; Yu & Shim, 2017). Building on this work, our experimental design uniquely allowed us to compare the magnitudes of both types of behavioural retro-cue benefits within a single experiment, and to correlate their strengths across participants. While the item benefit was larger than feature benefit, both were both highly robust. They were each evident across both colour and orientation reports and in both recall accuracy and response initiation times. Moreover, we found strong correlations between the benefits that followed item and feature cues, and qualitatively similar neural modulations, which suggest that the two types of retro-cueing benefits may share similar cognitive operations and resources.

The notion that both retro-cueing types yield behavioural benefits that are qualitatively similar was further supported by the similar retro-cueing effects on guess-rate, and target-response rate parameters estimated by the mixture model. At the same time, we observed that only item cues significantly enhanced precision and reduced the probability of swaps (non-target responses) – the latter being the only parameter that also differed significantly between item and feature retro-cue benefits. This difference is likely explained by the fact that swaps are calculated between items (not between features). Provided that feature-cues always concerned one feature, shared across both items, they may have helped up-regulate the relevant feature-dimension, but not to separate the two spatially-segregated items and thereby to reduce swap rates (in contrast to item cues that directly targeted the relevant item from the two memorised items).

In a strict account in which the primary unit of VWM is integrated items (Luck & Vogel, 1997; Vogel et al., 2001), one may predict that attention in VWM will primarily operate at the level of items, leaving little room for attentional facilitation of specific features that are shared among items. Alternatively, if VWM consists of a hierarchy of representations, with both item-level and feature-level representations (Bays, Wu, & Husain, 2011; Fougnie & Alvarez, 2011; Töllner, Conci, Müller, & Mazza, 2016; Töllner, Mink, & Müller, 2015); then one may expect that attention can operate similarly at distinct levels, depending on the nature of the task at hand. Our data are in line with a mixture of both scenarios – showing that attention can operate qualitatively similarly at both item and feature levels, while also revealing an additional benefit when attention is directed at two features of a single item (following item cues), compared to a single feature across two items (following feature cues).

At the same time, we note that attentional benefits in behavioural performance in VWM tasks need not only reflect changes in the quality of representational information. Factors related to prospective task preparation may also contribute (Myers, Stokes, & Nobre, 2017). Therefore, while our data provide clear evidence for the benefit of feature retro-cues – which is qualitatively similar to, and correlated with, the benefit following item cues – it remains possible that at least part of these benefits are due to factors other than a change in the underlying mnemonic representation (and this holds for both item and feature retro-cueing benefits).

In addition to the behavioural performance data, we also observed commonalities in the neural modulation following item and feature cues; both cases showing robust alpha attenuation over posterior electrodes, arising around the same time, with a similar magnitude. The neural responses therefore provide important relevant complementary data to our behavioural performance data. They provide more direct evidence for an early modulation in posterior (putatively visual) brain areas following both types of retro-cues; compatible with a modulation at the level of the memorised visual representations. However, because we used visual retro-cues, we cannot fully rule out the possibility that at least part of this modulation may be driven by differential visual processing of informative vs. a neutral retro-cues per se – though we note how our neutral retro-cues were designed to be similar to our informative retro-cues, ruling out more obvious differences due to bottom up visual features such as retro-cue size and saliency.

In conclusion, retro-cueing studies have typically shown that internally directed attention can prioritise a subset of mnemonic representations (Griffin & Nobre, 2003; Rerko, Souza, & Oberauer, 2014; Van Moorselaar, Olivers, Theeuwes, Lamme, Victor, & Sligte, 2015). These representations are typically thought of as integrated item of features bound together into a discrete mnemonic item (Luck & Vogel, 1997; Vogel et al., 2001). Our results show that attention can also effectively be directed to specific visual features that are shared across multiple items in memory – and for the first time reveal that such feature cues yield qualitatively similar (albeit weaker) behavioural benefits and neural modulations or latency, as do item cues, and that item and feature cueing benefits are correlated across individuals. We argue that retro-cues help place memorised visual stimuli into a goal-oriented format, such that relevant information at both the item and the feature-level can be optimised for upcoming task performance.

## Supplementary materials

**Supplementary Figure 1.**
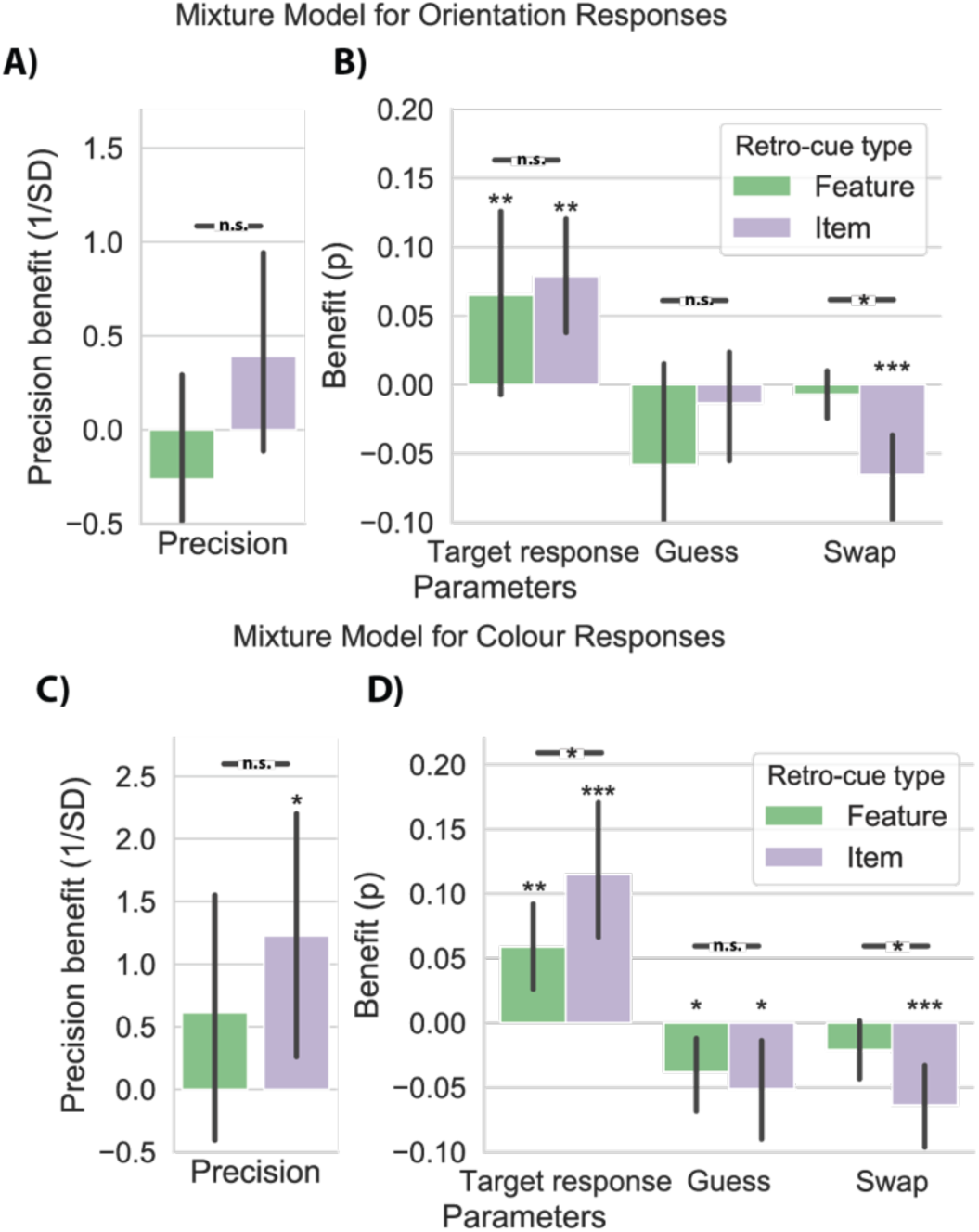
Mixture modelling parameters for colour and orientation. Mixture-model estimates for the benefit of orientation recall in A-B) and colour recall in C-D). Mixture model parameters include precision (A, C), target response, guess rate, and swap rate (B, D) for trials where item retro-cues or feature retro-cues were presented compared to neutral trials. Black asterisks indicate significant differences between item – and feature retro-cueing benefits. Asterisks above bars represent significance of a two-sided t-test of the model parameter benefit against zero. Error bars indicate 95% confidence intervals. * p < .05, ** p < .01, *** p < .001.

**Supplementary Figure 2.**
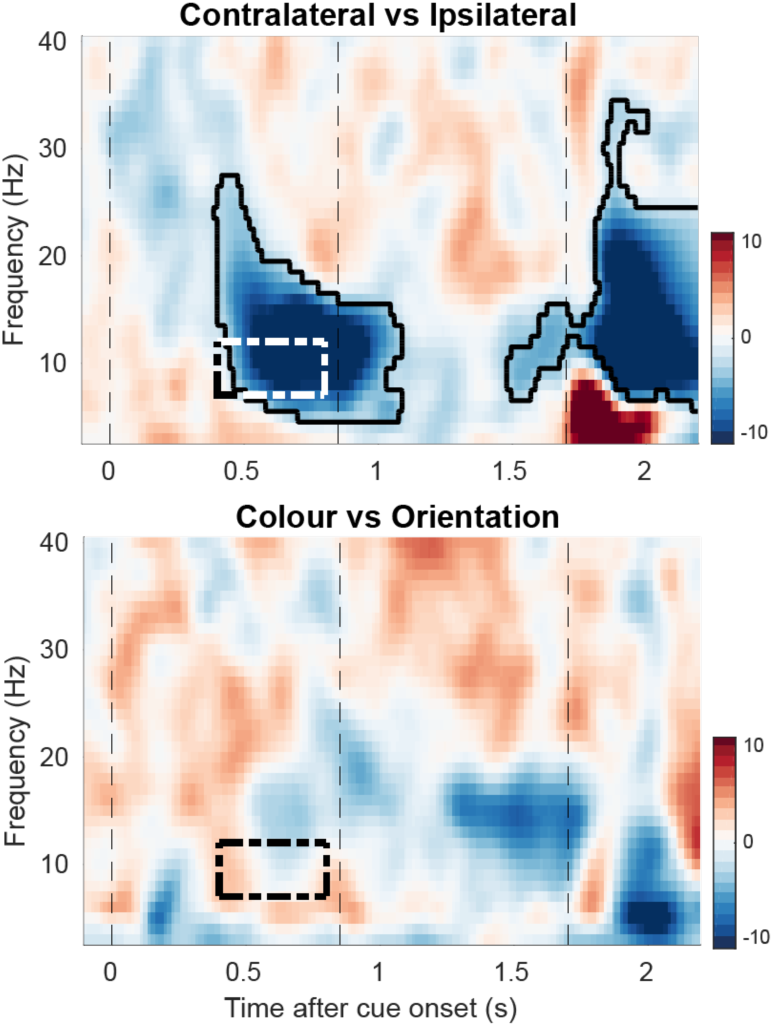
Induced neural differences between left vs. right item cues and between colour and orientation feature cues. Time-frequency maps associated with differences for informative item retro-cues directed at the contralateral vs. ipsilateral item (in selected left and right posterior electrodes; top) and for feature cues directed at colour vs orientation (in all posterior electrodes; bottom). Highlighted areas with the black solid outline indicate significant difference (permutation test, n = 30, cluster-forming threshold p < .05, corrected significance threshold p < .05). Dashed boxes indicate the window for which the topographies are depicted in **Figure 4**.

**Supplementary Figure 3.**
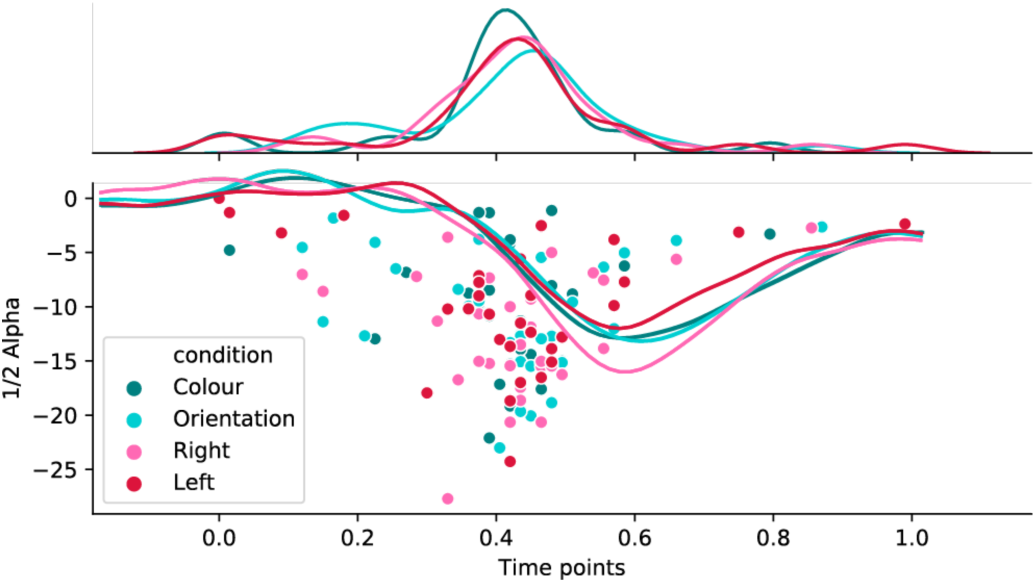
Onset times of the alpha attenuation after the retro-cue. The first time point where the alpha power (neutral – informative cue) reached half of its minimum value in the interval from 0 to 1000 ms after the retro-cue onset was taken as the alpha attenuation latency. The dots represent alpha attenuation latency times for individual subjects and different cueing conditions with a density plot on the top showing the density of the dots along the x-axis for each condition. The line plot illustrates the average alpha power for each condition after subtracting their relative neutral retro-cues.

**Supplementary table 1.** Main effects and interactions for the congruence or incongruence of the distractor with the probed memory feature, tested with a 2 × 2 × 2 rmANOVA with the factors distractor congruence, cue informativeness, and block type, separately for each of our four dependent variables.

**B) Within Subjects Effects for Orientation recall: Error**
| | Sum of Squares | df | Mean Square | F | p | $\eta^2$ |
| --- | --- | --- | --- | --- | --- | --- |
| Distractor | 0.022 | 1 | 0.022 | 5.515 | 0.026 | 0.160 |
| Residual | 0.116 | 29 | 0.004 |  |  |  |
| Retro-cue type * Distractor | 7.879e -4 | 1 | 7.879e -4 | 0.256 | 0.617 | 0.009 |
| Residual | 0.089 | 29 | 0.003 |  |  |  |
| Informativeness * Distractor | 4.938e -4 | 1 | 4.938e -4 | 0.142 | 0.709 | 0.005 |
| Residual | 0.101 | 29 | 0.003 |  |  |  |
| Retro-cue type * Informativeness * Distractor | 5.667e -5 | 1 | 5.667e -5 | 0.026 | 0.872 | 0.001 |
| Residual | 0.063 | 29 | 0.002 |  |  |  |

**B) Within Subjects Effects for Colour recall: Error**
| | Sum of Squares | df | Mean Square | F | p | $\eta^2$ |
| --- | --- | --- | --- | --- | --- | --- |
| Distractor | 7.333e -4 | 1 | 7.333e -4 | 0.286 | 0.597 | 0.010 |
| Residual | 0.074 | 29 | 0.003 |  |  |  |
| Retro-cue type * Distractor | 0.001 | 1 | 0.001 | 1.290 | 0.265 | 0.043 |
| Residual | 0.023 | 29 | 7.788e -4 |  |  |  |
| Informativeness * Distractor | 0.001 | 1 | 0.001 | 0.643 | 0.429 | 0.022 |
| Residual | 0.050 | 29 | 0.002 |  |  |  |
| Retro-cue type * Informativeness * Distractor | 2.928e -4 | 1 | 2.928e -4 | 0.295 | 0.591 | 0.010 |
| Residual | 0.029 | 29 | 9.912e -4 |  |  |  |

**C) Within Subjects Effects for Orientation recall: RT**
| | Sum of Squares | df | Mean Square | F | p | $\eta^2$ |
| --- | --- | --- | --- | --- | --- | --- |
| Distractor | 8427 | 1 | 8427 | 1.939 | 0.174 | 0.063 |
| Residual | 126029 | 29 | 4346 |  |  |  |
| Retro-cue type * Distractor | 6784 | 1 | 6784 | 2.556 | 0.121 | 0.081 |
| Residual | 76971 | 29 | 2654 |  |  |  |
| Informativeness * Distractor | 1807 | 1 | 1807 | 0.549 | 0.465 | 0.019 |
| Residual | 95526 | 29 | 3294 |  |  |  |
| Retro-cue type * Informativeness * Distractor | 2004 | 1 | 2004 | 0.652 | 0.426 | 0.022 |
| Residual | 89135 | 29 | 3074 |  |  |  |

**D) Within Subjects Effects for Colour recall: RT**
| | Sum of Squares | df | Mean Square | F | p | $\eta^2$ |
| --- | --- | --- | --- | --- | --- | --- |
| Distractor | 6517.18 | 1 | 6517.18 | 0.629 | 0.434 | 0.021 |
| Residual | 300276.91 | 29 | 10354.38 |  |  |  |
| Retro-cue type * Distractor | 339.93 | 1 | 339.93 | 0.119 | 0.733 | 0.004 |
| Residual | 83020.69 | 29 | 2862.78 |  |  |  |
| Informativeness * Distractor | 1762.09 | 1 | 1762.09 | 0.698 | 0.410 | 0.024 |
| Residual | 73178.64 | 29 | 2523.40 |  |  |  |
| Retro-cue type * Informativeness * Distractor | 54.09 | 1 | 54.09 | 0.023 | 0.879 | 0.001 |
| Residual | 66963.22 | 29 | 2309.08 |  |  |  |

## References

Alvarez, G. A., & Cavanagh, P. (2004). The Capacity of Visual Short-Term Memory Is Set Both by Visual Information Load and by Number of Objects, 15(2), 106–111.

Baddeley, A. D. (1992). Working Memory. Science, 255(Jan), 556–559. https://doi.org/DOI:10.1126/science.1736359

Baddeley, A. D. (2003). Working memory: looking back and looking forward. Nature Reviews. Neuroscience, 4(10), 829–839. https://doi.org/10.1038/nrn1201

Bays, P. M., Catalao, R. F. G., & Husain, M. (2009). The precision of visual working memory is set by allocation of a shared resource. Journal of Vision, 9(10), 7.1-11. https://doi.org/10.1167/9.10.7

Bays, P. M., & Husain, M. (2008). Dynamic shifts of limited working memory resources in human vision. Science (New York, N.Y.), 321(5890), 851–854. https://doi.org/10.1126/science.1158023

Bays, P. M., Wu, E. Y., & Husain, M. (2011). Storage and binding of object features in visual working memory. Neuropsychologia, 49(6), 1622–1631. https://doi.org/10.1016/j.neuropsychologia.2010.12.023

Brainard, D. H. (1997). The Psychophysics Toolbox. Spatial Vision, 10, 433–436. https://doi.org/10.1163/156856897X00357

Fougnie, D., & Alvarez, G. A. (2011). Object features fail independently in visual working memory: Evidence for a probabilistic feature-store model. Journal of Vision, 11(12), 3–3. https://doi.org/10.1167/11.12.3.Introduction

Griffin, I. C., & Nobre, A. C. (2003). Orienting attention to locations in internal representations. Journal of Cognitive Neuroscience, 15(8), 1176–1194. https://doi.org/10.1162/089892903322598139

Hyvärinen, A. (1999). Fast and robust fixed-point algorithms for independent component analysis. IEEE Transactions on Neural Networks, 10(3), 626–634. https://doi.org/10.1109/72.761722

Kuo, B.-C., Stokes, M. G., & Nobre, A. C. (2011). Attention Modulates Maintenance of Representations in Visual Short-term Memory. Journal of Cognitive Neuroscience, 24(1), 51–60. https://doi.org/10.1162/jocn_a_00087

Landman, R., Spekreijse, H., & Lamme, V. A. F. (2003). ScienceDirect - Vision Research : Large capacity storage of integrated objects before change blindness. Vision Research, 43, 149–164.

Luck, S. J., & Vogel, E. K. (1997). The capacity of visual working memory for features and conjunctions. Nature, 390(6657), 279–281. https://doi.org/10.1038/36846

Maris, E., & Oostenveld, R. (2007). Nonparametric statistical testing of EEG- and MEG-data. Journal of Neuroscience Methods, 164(1), 177–190. https://doi.org/10.1016/j.jneumeth.2007.03.024

Myers, N. E., Stokes, M. G., & Nobre, A. C. (2017). Prioritizing Information during Working Memory: Beyond Sustained Internal Attention. Trends in Cognitive Sciences, 21(6), 449–461. https://doi.org/10.1016/j.tics.2017.03.010

Myers, N. E., Walther, L., Wallis, G., Stokes, M. G., & Nobre, A. C. (2015). Temporal Dynamics of Attention during Encoding versus Maintenance of Working Memory: Complementary Views from Event-related Potentials and Alpha-band Oscillations. Journal of Cognitive Neuroscience, 27(3), 492–508. https://doi.org/10.1162/jocn

Niklaus, M., Nobre, A. C., & Van Ede, F. (2017). Feature-based attentional weighting and spreading in visual working memory. Scientific Reports, 7, 1–10. https://doi.org/10.1038/srep42384

Nobre, A. C. (2018). Sensation, Perception, and Action. In J. T. Wixed & J. T. Serences (Eds.), Stevens’ Handbook of Experimental Psychology and Cognitive Neuroscience (Fourth Edi, pp. 241–316). New York: John WIley & Sons Incl.

Oostenveld, R., Fries, P., Maris, E., & Schoffelen, J. M. (2011). FieldTrip: Open source software for advanced analysis of MEG, EEG, and invasive electrophysiological data. Computational Intelligence and Neuroscience, 2011. https://doi.org/10.1155/2011/156869

Park, Y. E., Sy, J. L., Hong, S. W., & Tong, F. (2017). Reprioritization of Features of Multidimensional Objects Stored in Visual Working Memory. Psychological Science, 28(12), 1773–1785. https://doi.org/10.1177/0956797617719949

Pilling, M., & Barrett, D. J. K. (2016). Dimension-based attention in visual short-term memory. Memory and Cognition, 44(5), 740–749. https://doi.org/10.3758/s13421-016-0599-6

Poch, C., Capilla, A., Hinojosa, J. A., & Campo, P. (2017). Selection within working memory based on a color retro-cue modulates alpha oscillations. Neuropsychologia, 106(July), 133–137. https://doi.org/10.1016/j.neuropsychologia.2017.09.027

Rerko, L., Souza, A. S., & Oberauer, K. (2014). Retro-cue benefits in working memory without sustained focal attention. Memory and Cognition, 42(5), 712–728. https://doi.org/10.3758/s13421-013-0392-8

Rosner, B. (1983). Percentage Outlier Points for Generalized ESD Many-Procedure. Technometrics, 25(2), 165–172.

Souza, A. S., & Oberauer, K. (2016). In search of the focus of attention in working memory: 13 years of the retro-cue effect. Attention, Perception, and Psychophysics, 78(7), 1839–1860. https://doi.org/10.3758/s13414-016-1108-5

Töllner, T., Conci, M., Müller, H. J., & Mazza, V. (2016). Attending to multiple objects relies on both feature- and dimension-based control mechanisms: Evidence from human electrophysiology. Attention, Perception, and Psychophysics, 78(7), 2079–2089. https://doi.org/10.3758/s13414-016-1152-1

Töllner, T., Mink, M., & Müller, H. J. (2015). Searching for targets in visual working memory: Investigating a dimensional feature bundle (DFB) model. Annals of the New York Academy of Sciences, 1339(1), 32–44. https://doi.org/10.1111/nyas.12703

van Ede, F. (2018). Mnemonic and attentional roles for states of attenuated alpha oscillations in perceptual working memory: a review. European Journal of Neuroscience, 48(7), 2509–2515. https://doi.org/10.1111/ejn.13759

van Ede, F., Niklaus, M., & Nobre, A. C. (2017). Temporal Expectations Guide Dynamic Prioritization in Visual Working Memory through Attenuated α Oscillations. The Journal of Neuroscience, 37(2), 437–445. https://doi.org/10.1523/JNEUROSCI.2272-16.2017

Van Moorselaar, D., Olivers, C. N. L., Theeuwes, J., Lamme Victor, A. F., & Sligte, I. G. (2015). Forgotten But Not Gone : Retro-Cue Costs and Benefits in a Double-Cueing Paradigm Suggest Multiple States in Visual … Journal of Experimental Psychology : Learning, Memory, and Cognition. Journal of Experimental Psychology, 41(6), 1755–1763. https://doi.org/10.1037/xlm0000124

Vogel, E. K., McCollough, A. W., & Machizawa, M. G. (2005). Neural measures reveal individual differences in controlling access to working memory. Nature, 438(7067), 500–503. https://doi.org/10.1038/nature04171

Vogel, E. K., Woodman, G. F., & Luck, S. J. (2001). Storage of features, conjunctions, and objects in visual working memory. Journal of Experimental Psychology, 27(1), 92–114.

Wallis, G., Stokes, M. G., Cousijn, H., Woolrich, M., & Nobre, A. C. (2015). Frontoparietal and Cingulo-opercular Networks Play Dissociable Roles in Control of Working Memory. Journal of Cognitive Neuroscience, 27(10), 2019–2034. https://doi.org/https://doi.org/10.1162/jocn_a_00838

Wolff, M. J., Jochim, J., Akyürek, E. G., & Stokes, M. G. (2017). Dynamic hidden states underlying working-memory-guided behavior. Nature Neuroscience, 20(6), 864–871. https://doi.org/10.1038/nn.4546

Ye, C., Hu, Z., Ristaniemi, T., Gendron, M., & Liu, Q. (2016). Retro-dimension-cue benefit in visual working memory. Scientific Reports, 6, 1–13. https://doi.org/10.1038/srep35573

Yu, Q., & Shim, W. M. (2017). Occipital, parietal, and frontal cortices selectively maintain task-relevant features of multi-feature objects in visual working memory. NeuroImage, 157(May), 97–107. https://doi.org/10.1016/j.neuroimage.2017.05.055

Zhang, W., & Luck, S. J. (2008). Discrete fixed-resolution representations in visual working memory. Nature, 453(7192), 233–235. https://doi.org/10.1038/nature06860

